# Impact of Sample Type and DNA Isolation Procedure on Genomic Inference of Microbiome Composition

**DOI:** 10.1101/064394

**Authors:** Berith E. Knudsen, Lasse Bergmark, Patrick Munk, Oksana Lukjancenko, Anders Priemé, Frank M. Aarestrup, Sünje J. Pamp

## Abstract

Explorations of complex microbiomes using genomics greatly enhance our understanding about their diversity, biogeography, and function. The isolation of DNA from microbiome specimens is a key prerequisite for such examinations, but challenges remain in obtaining sufficient DNA quantities required for certain sequencing approaches, achieving accurate genomic inference of microbiome composition, and facilitating comparability of findings across specimen types and sequencing projects. These aspects are particularly relevant for the genomics-based global surveillance of infectious agents and antimicrobial resistance from different reservoirs. Here, we compare in a stepwise approach a total of eight commercially available DNA extraction kits and 16 procedures based on these for three specimen types (human feces, pig feces, and hospital sewage). We assess DNA extraction using spike-in controls, and different types of beads for bead-beating facilitating cell lysis. We evaluate DNA concentration, purity, and stability, and microbial community composition using 16S rRNA gene sequencing and for selected samples using shotgun metagenomic sequencing. Our results suggest that inferred community composition was dependent on inherent specimen properties as well as DNA extraction method. We further show that bead-beating or enzymatic treatment can increase the extraction of DNA from Gram-positive bacteria. Final DNA quantities could be increased by isolating DNA from a larger volume of cell lysate compared to standard protocols. Based on this insight, we designed an improved DNA isolation procedure optimized for microbiome genomics that can be used for the three examined specimen types and potentially also for other biological specimens. A standard operating procedure is available from: https://dx.doi.org/10.6084/m9.figshare.3475406.

**IMPORTANCE:** Sequencing-based analyses of microbiomes may lead to a breakthrough in our understanding of the microbial world associate with humans, animals, and the environment. Such insight could further the development of innovative ecosystem management approaches for the protection of our natural resources, and the design of more effective and sustainable solutions to prevent and control infectious diseases. Genome sequence information is an organism- (pathogen-) independent language that can be used across sectors, space, and time. Harmonized standards, protocols, and workflows for sample processing and analysis can facilitate the generation of such actionable information. In this study, we assessed several procedures for the isolation of DNA for next-generation sequencing. Our study highlights several important aspects to consider in the design and conduction of sequence-based analysis of microbiomes. We provide a standard operating procedure for the isolation of DNA from a range of biological specimens particularly relevant in clinical diagnostics and epidemiology.

## INTRODUCTION

Microbial communities fulfill central roles in biological systems, such as in human, animal, and environmental ecosystems. Genomics-based interrogations of these communities can provide unprecedented insight into their composition and function, and reveal general principles and rules about their ecology and evolution (1–4). Genomics-based microbiome analyses can also have important practical implications, such as for the diagnosis and management of infectious diseases. Together with relevant metadata, attribute data, and appropriate bioinformatics and statistical approaches, genomic sequencing data could enable the global surveillance of emerging and re-emerging infectious diseases, and teach us about the reservoirs and transmission pathways of pathogens (5–7). Ultimately, genomics-based information about infectious disease epidemiology may help us to predict, prevent, and control infectious diseases faster, more precisely, and more sustainably.

In order to facilitate large-scale microbiome analyses, harmonized standards for sample handling and data analysis need to be ensured. To be able to establish pathogen reservoirs and transmission pathways, specimens from different sources, such as from humans, animals, and the environment, will need to be examined. For genomics analysis, the DNA needs to be isolated from the specimens for DNA sequencing. However, DNA isolation methods are often only evaluated and established in the context of specimens from an individual source (e.g. human fecal specimens), and seldom across a variety of specimen types (8–12), which is addressed in the present study. Current sequencing technologies, such as Illumina MiSeq and Hiseq, PacBio, IonTorrent, and Nanopore systems, still require large initial DNA template quantities, particularly from the perspective of PCR-free metagenomics-based analysis. In contrast, 16S rRNA gene profiling can reveal a bacterial and archaeal composition for samples with low initial DNA template quantities. In metagenomics, low quantities of input DNA can result in low sequencing data output, and impact the inferred microbial community composition (13). Hence, modified DNA isolation protocols for increasing DNA quantities obtained from different types of specimens are desirable.

Here, we examine three specimen types (human feces, animal feces, and sewage), a total of eight commercially available DNA isolation kits, and a number of protocol modifications in regard to output DNA (quantity, purity, stability) and microbiome composition (16S rRNA gene profiling, metagenomics). Our results suggest that both, the specimen itself as well as the DNA isolation procedure, can affect DNA quantity and quality, and inferred microbiome composition. Based on the insight gained, we have developed an improved laboratory protocol that can be used for DNA isolations from a variety of biological specimens.

## RESULTS

### DNA concentration, purity, and stability depend on the type of specimen and DNA isolation method

We extracted DNA from human feces, pig feces, and hospital sewage, using seven commonly used DNA isolation kits and determined DNA concentration, purity, and stability of the isolated DNA (Fig. 1A and Table 1). The DNA concentrations varied greatly (Fig. 1B; see also Table S1A in the supplemental material). For human feces, the highest DNA concentrations were obtained using the EasyDNA, MagNAPure, and QIAStool procedure, for pig feces using the EasyDNA, QIAStool, and QIAStool+BB procedures, and for sewage using the MagNAPure and EasyDNA procedure, while for three methods the DNA concentration from sewage was below the detection limit. On average across the three types of specimen, the highest DNA concentrations were obtained using EasyDNA (44.96 ng/μl +/ࢤ 20.99 SEM) and QIAStool 27.88 ng/μl +/− 2.55 SEM), and the lowest using the PowerSoil.HMP (1.55 ng/μl +/−0.31 SEM) and InnuPURE (7.77 ng/μl +/− 5.54 SEM) methods.

**FIG 1.**
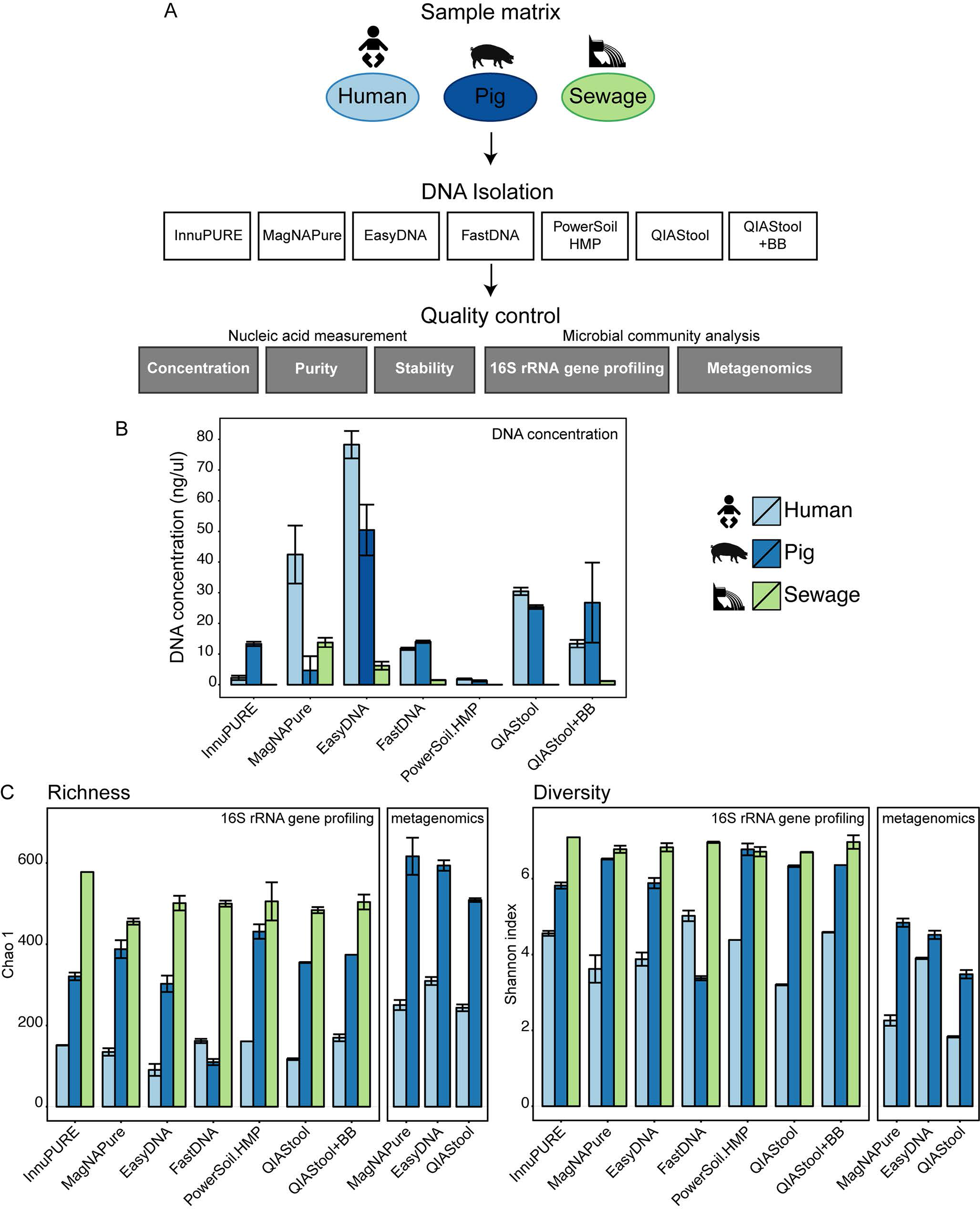
Comparison of DNA extraction methods. (A) Experimental design. Human feces, pig feces, and hospital sewage were extracted using seven different DNA extraction methods (see also Table 1): InnuPure^®^ C16, MagNA Pure LC DNA isolation Kit III, Easy-DNA™ gDNA Purification Kit, MP FastDNA™ Spin Kit, PowerSoil^®^ DNA Isolation kit, QIAamp^®^ DNA Stool Mini Kit, QIAamp^®^ DNA Stool Mini Kit + Bead Beating (For details see Materials and Methods). DNA concentration, purity, and stability were examined, and microbial community composition determined using 16S rRNA gene profiling and metagenomics (selected samples). (B) DNA from each method was dissolved in 100 ul solution and DNA concentrations were determined using Qubit^®^ dsDNA BR Assay Kit measurements. Values represent averages from duplicate or triplicate DNA extractions (See also Supplemental Table S1A). (C) Ecological richness (Chao 1) and diversity (Shannon index) were determined based on contingency tables from 16S rRNA gene profiling and metagenomic sequencing data at OTU and species levels, respectively (See also Supplemental Table S1B).

**TABLE 1.**
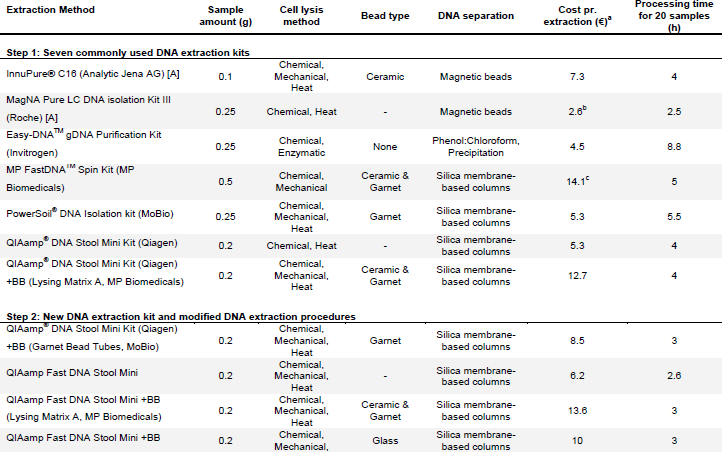
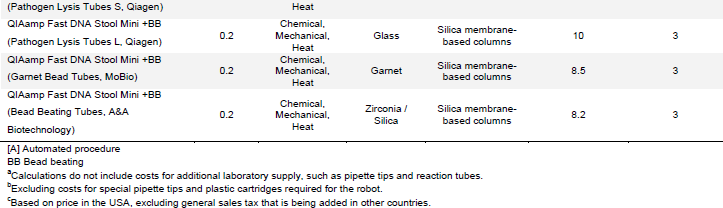
Overview of DNA extraction procedures

With regard to DNA purity, the best results for human and pig feces were obtained using the EasyDNA, QIAStool, and QIAStool+BB procedure (see Table S1A in the supplemental material). The DNA was generally stable for at least 7 days when stored at room temperature (22°C) with some exceptions (see Table S1A in the supplemental material). A decrease in DNA concentration over time was observed for example for the human feces when extracted with EasyDNA (57% decease in DNA concentration) or MagNAPure (21% decrease in DNA concentration), suggesting the presence of DNases in these extracts. In some cases, an increase in DNA concentration over time was observed, such as for the pig feces when extracted with EasyDNA (32% increase in DNA concentration). An increase in DNA concentration over time at room temperature was previously shown to be related to the hyperchromicity of DNA, and dependent on the DNA concentration and ionic strength of the solution (14).

### Microbial richness and diversity are influenced by DNA isolation procedure

For the human fecal specimen, the highest bacterial Operational Taxonomic Unit (OTU) richness and diversity were detected using the QIAStool+BB and FastDNA methods, followed by InnuPURE and PowerSoil.HMP as assessed by 16S rRNA gene profiling (Fig. 1C; see also Table S1B in the supplemental material). In comparison, the determined richness and diversity for the EasyDNA method was low, and the relative abundance of Ruminococcaceae and Bifidobacteriaceae dominated the composition compared to the extracts from the other methods (Fig. 1C; see also Fig. S1A in the supplemental material). Thirty-nine samples (human feces, pig feces, and sewage) with high DNA concentration were selected and examined using metagenomic sequencing. In this assessment, the species richness and diversity for human feces was highest for the EasyDNA procedure, and a high relative abundance of Ruminococcaceae and Bifidobacteriaceae was apparent in this analysis as well (see Fig. S1A in the supplemental material).

For the pig fecal specimen, the highest bacterial richness and diversity were detected using the PowerSoil.HMP and MagNAPure methods, followed by QIAStool+BB (Fig. 1C; see also Table S1B in the supplemental material). Similarly, richness and diversity were highest using the MagNAPure and EasyDNA methods when assessed using metagenomics. Based on 16S rRNA gene profiling, the richness and diversity for the FastDNA method were lower compared to all other methods, and the relative abundance of Clostridiaceae and Turicibacteraceae was higher and the abundance of Prevotellaceae and Ruminococcaceae lower using this method, compared to the other methods (Fig. 1C; see also Fig. S1A in the supplemental material).

For the sewage specimen, the highest bacterial richness and diversity was detected using the InnuPURE method, followed by PowerSoil.HMP and QIAStool+BB, and similar levels were achieved using the other methods (Fig. 1C; see also Table S1B in the supplemental material). The relative abundance of Clostridiaceae was highest in the samples extracted using EasyDNA, and the abundance of Enterobacteriales highest in the samples extracted using PowerSoil.HMP.

Overall, the relative abundance of predicted Gram-positive bacteria was highest in the human and sewage specimens when extracted with the EasyDNA method, and highest in the pig specimen when extracted using the FastDNA method (see Fig. S2 in the supplemental material). The abundance of predicted Gram-positive bacteria was lowest using MagNAPure and QIAStool, the two methods that did neither include a bead-beating step nor specific enzymatic cell-wall digestion.

### Microbial community composition depends on the choice of DNA isolation procedure

The microbial communities from the three types of specimen clustered separately according to specimen type when examined in PCoA Bray-Curtis ordination, and not according to DNA isolation procedure (see Fig. S3 in the supplemental material), indicating that the largest differences between these samples are driven by the inherent microbiota composition. Bray-Curtis dissimilarity distance analysis carried out separately for each of the three specimens revealed that the samples largely clustered according to DNA isolation procedure (Fig. 2A-C). For the human fecal specimen, the bacterial community composition derived from the EasyDNA isolation differed from the communities obtained using all other methods (Fig. 2A), which is in agreement with the observations on microbial richness (above). The Bray-Curtis distances between the samples from InnuPURE, MagNAPure, FastDNA, PowerSoil.HMP, QIAStool, and QIAStool+BB DNA isolations were on average 0.337 +/−0.012 SEM, whereas the distances between these and the ones derived from the EasyDNA procedure were on average 0.825 +/− 0.014 SEM.

**FIG 2.**
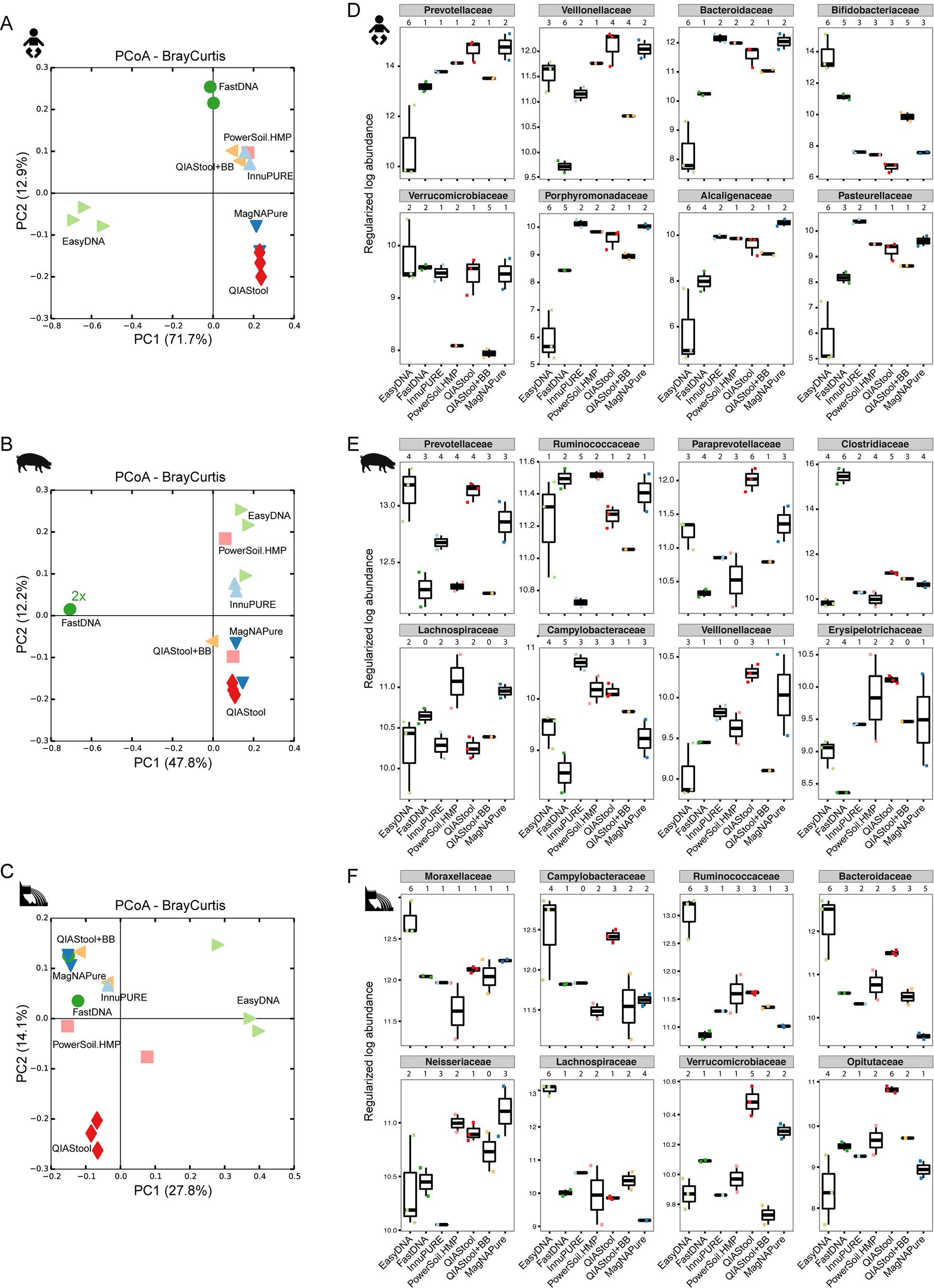
Microbial community dissimilarity. The dissimilarity between the microbiotas from the human, pig, and sewage samples based on DNA extraction methods was examined using Principal Coordinates Analysis of Bray-Curtis distances (A-C) and differential abundance analysis using DESeq2 (D-F) from 16S rRNA amplicon data. (A-F) For the PCoA Bray-Curtis ordination analysis only samples with 800 or more reads were included. (D-F) For the differential abundance analysis pairwise testing by DNA extraction method was performed, and bacterial families were considered significantly differentially abundant if their adjusted P-value was <0.1 (see also Table S2 in the supplemental material). Examples for differentially abundant families are shown that are among the top10 most abundant taxa found in the sample, respectively. For each family, the total number of DNA isolation procedures, that exhibit significantly different abundance values compared to a particular DNA isolation procedure, are indicated above the plot, respectively.

For the pig fecal specimen, the bacterial communities derived from the FastDNA isolation differed from all other communities (Fig. 2B). The average Bray-Curtis distance between the samples originating from all but the FastDNA procedure was on average 0.473 +/− 0.008 SEM, whereas the distance between these and the ones derived from the FastDNA procedure was on average 0.877 +/− 0.007 SEM.

For the hospital sewage specimen, the bacterial communities originating from the EasyDNA method differed from all others (average Bray-Curtis distance 0.600 +/− 0.006 SEM) (Fig. 2C), similar to the human fecal matrix (Fig. 2A). In addition, the communities originating from the QIAStool DNA isolation differed from all others (average Bray-Curtis distance 0.514 +/− 0.009 SEM), whereas the average Bray-Curtis distance between all but the QIAStool and EasyDNA samples was 0.460 +/−0.11 SEM on average.

### Distinct taxa account for the differences observed between DNA isolation methods

To quantify the effect of DNA isolation method on microbial community composition we tested for differential abundance of taxa between the communities derived from the different DNA isolation methods using DESeq2 analyses. In pairwise comparisons, significant differences between the DNA isolation methods were observed (Fig. 2D-F; see also Table S2 in the supplemental material).

The most abundant family on average in the human fecal specimen was Prevotellaceae (Bacteroidetes), and its abundance was significantly lower in the samples extracted with EasyDNA as compared to all other methods (e.g. 18.3-fold lower in EasyDNA vs.

QIAStool, adjusted p-value 1.91^−6^) (Fig. 2D; see also Table S2 in the supplemental material). Similarly, the abundance of Bacteroidaceae (Bacteroidetes), Porphyromonadaceae (Bacteroidetes), Alcaligenaceae (β-Proteobacteria), and Pasteurellaceae (γ-Proteobacteria) was lower in the samples from the EasyDNA isolation compared to the other methods. In contrast, the abundance of Bifidobacteriaceae (Actinobacteria) was higher in the samples originating from the EasyDNA procedure compared to all other methods (e.g. 770-fold higher in EasyDNA vs. QIAStool, adjusted p-value 7.49^−57^). The abundance of Verrucomicrobiaceae (Verrucomicrobia) was significantly lower in the samples from the QIAStool+BB and PowerSoil.HMP DNA isolations (e.g. 4.15-fold lower in QIAStool+BB vs. QIAStool, adjusted p-value 0.001).

The most abundant family on average in the pig fecal specimen was Prevotellaceae (Bacteroidetes), and its abundance differed significantly between the DNA isolation procedures (e.g. 2.3-fold lower in EasyDNA vs. PowerSoil.HMP, adjusted p-value 1.28^−5^) (Fig. 2E; see also Table S2 in the supplemental material). The abundance of Clostridiaceae (Clostridia), the on average fourth most abundant family in the pig feces, was significantly higher in the samples extracted by the FastDNA method (e.g. 166-fold higher in FastDNA vs. EasyDNA, adjusted p-value 7.35^−110^).

Moraxellaceae (γ-Proteobacteria) was the most abundant family on average in the hospital sewage, and its abundance was significantly higher in the samples from the EasyDNA isolation compared to other DNA isolation methods (e.g. 2.6-fold higher in EasyDNA vs. PowerSoil.HMP, adjusted p-value 3.82^−5^) (Fig. 2F; see also Table S2 in the supplemental material). Ruminococcaceae (Clostridia), the on average third most abundant family in sewage, were also significantly more abundant in the samples from the EasyDNA isolation compared to other DNA isolation procedures (e.g. 7.3-fold higher in EasyDNA vs. FastDNA, adjusted p-value 4.28^−17^).

### DNA isolation procedure affects the abundance of taxa differently across specimens

Given that differential taxa abundances were observed for the different DNA isolation procedures for the three specimen types, we investigated whether the abundance differed in the same way between DNA isolation procedures across specimens. For example, we were asking: If taxon A is observed at a higher abundance upon DNA isolation with method X compared to method Y in specimen type 1, is this taxon also observed at a higher abundance upon DNA isolation with method X compared to method Y in specimen type 2? We examined taxa that were detected in all three specimen types, and selected representative families from different phyla (Fig. 3). Similar patterns of differential abundance were observed for certain taxa across specimen types, with exceptions, including two families from the Bacteroidetes phylum. The abundance of Prevotellaceae and Bacteroidaceae was significantly lower when human fecal specimen were extracted with EasyDNA compared to other methods. In contrast, these two families were observed at a significantly higher abundance when sewage was extracted with EasyDNA compared to other methods (Fig. 3).

**FIG 3.**
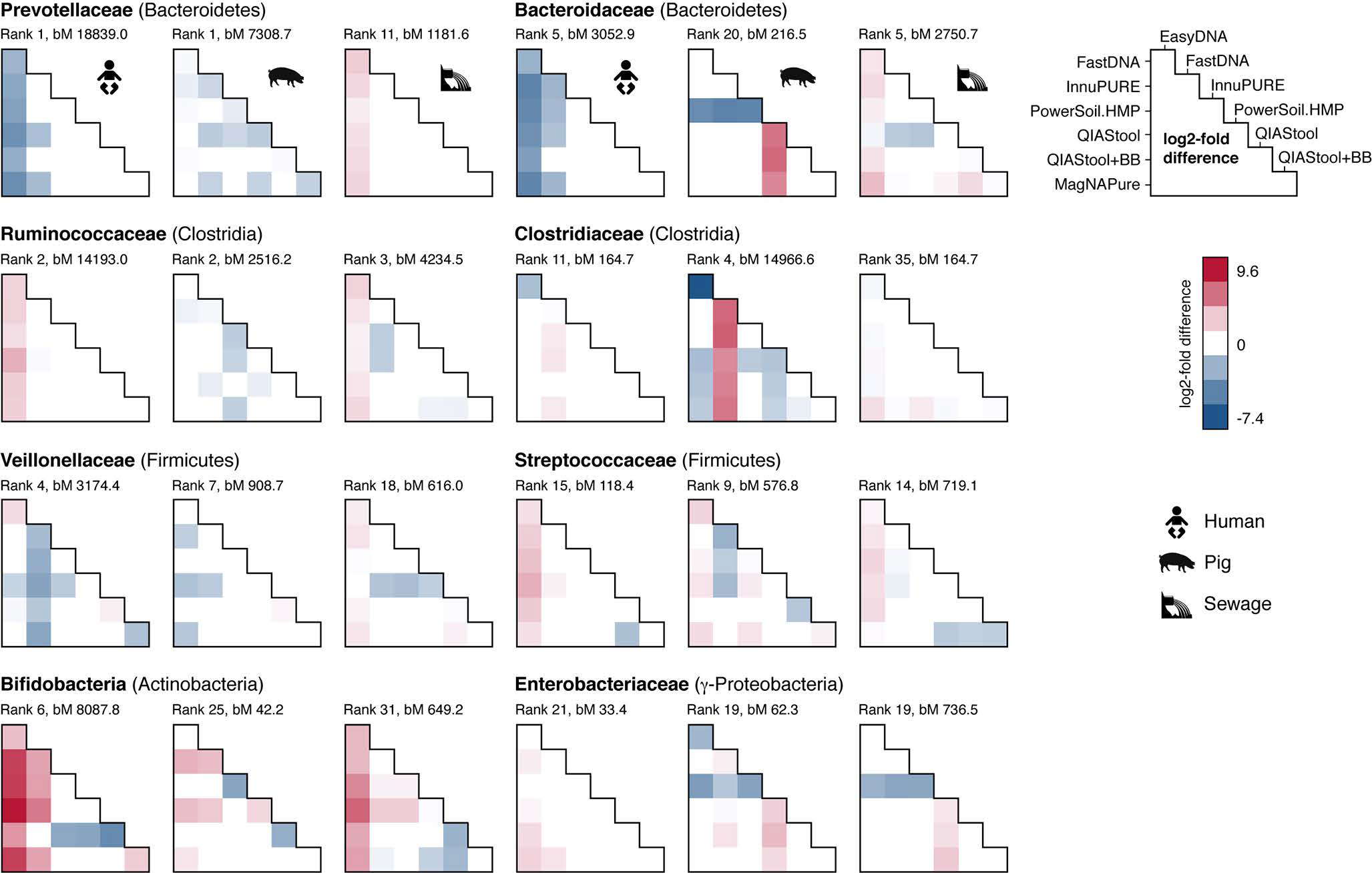
Differential abundance of bacterial families. Pairwise testing by DNA extraction method was performed using DESeq2, and the log2-fold difference displayed (column vs. rows) for selected families present in all sample matrices if their adjusted P-value was <0.1 (see also Table S2 in the supplemental material). The rank abundance position for each family per sample matrix type is noted according to their regularized log abundance. The baseMean (bM) indicates the mean of negative-binominal-based normalized read counts. The pairwise comparisons based on relative abundance normalization (total-sum scaling) of the bacterial families for the different DNA isolation procedures and three sample types is available though figshare at <u>https://dx.doi.org/10.6084/m9.figshare.3811254.</u>

Likewise, Ruminococcaceae of the phylum Clostridia were observed at a significantly higher abundance in human fecal and hospital sewage samples but not in pig fecal samples when extracted with the EasyDNA method compared to other methods. The same pattern was however not observed for all families of the phylum Clostridia. Clostridiaceae abundance appeared higher in human and pig feces when extracted with FastDNA compared to other methods, and Clostridiaceae abundance appeared higher in sewage when extracted using the EasyDNA method compared to other methods (Fig. 3).

Thus, we found significant differences in the abundance of certain families according to specimen type, which sometimes depend on the DNA isolation procedure. Some of the differential abundance patterns were similar across the three types of specimens, while others differed.

### Detection of spiked bacteria is dependent on DNA isolation procedure and specimen type

In order to quantify DNA isolation efficiency, we spiked the three specimen with known numbers of two bacterial representatives, namely *Salmonella enterica* serotype Typhimurium DT104 (Gram-negative) and *Staphylococcus aureus* ST398 (Gram-positive) in a CFU ratio of 1.02. Both, *S. enterica* and *S. aureus* were present in negligible numbers in the three specimens before spiking. DNA was isolated from these samples using the seven different DNA isolation methods, and the abundance of the two strains determined using 16S rRNA gene profiling, and for some samples also using metagenomics. Based on 16S rRNA gene profiling, the spiked organisms accounted for an average abundance of 1.0% (+/−0.29 SEM) Enterobacteriaceae, and 0.29% (+/−0.11 SEM) Staphylococcaceae across the three types of specimen.

Using QIAStool, a DNA isolation method that does not involve a bead-beating step, the abundance of Enterobacteriaceae was higher in the spiked human fecal specimen than expected, with an Enterobacteriaceae/Staphylococcaceae (E/S) ratio of 13.9 (Fig. 4A). This ratio was lower in the spiked human fecal specimen using InnuPURE, FastDNA, PowerSoil.HMP, and QIAStool+BB, which are all methods that involve a bead-beating step (E/S ratio range 0.3-2.3). The EasyDNA method involves an additional enzymatic lysis step, and using this method the determined E/S ratio was 3.7. Using the MagNAPure method no or lower read numbers assigned to Staphylococcaceae were detected in the spiked samples compared to not spiked samples in the human fecal specimen, and hence the ratio resulted in negative values (Fig. 4A). A similar result was obtained when the samples were examined using metagenomics (see Fig. S4 in the supplemental material).

**FIG 4.**
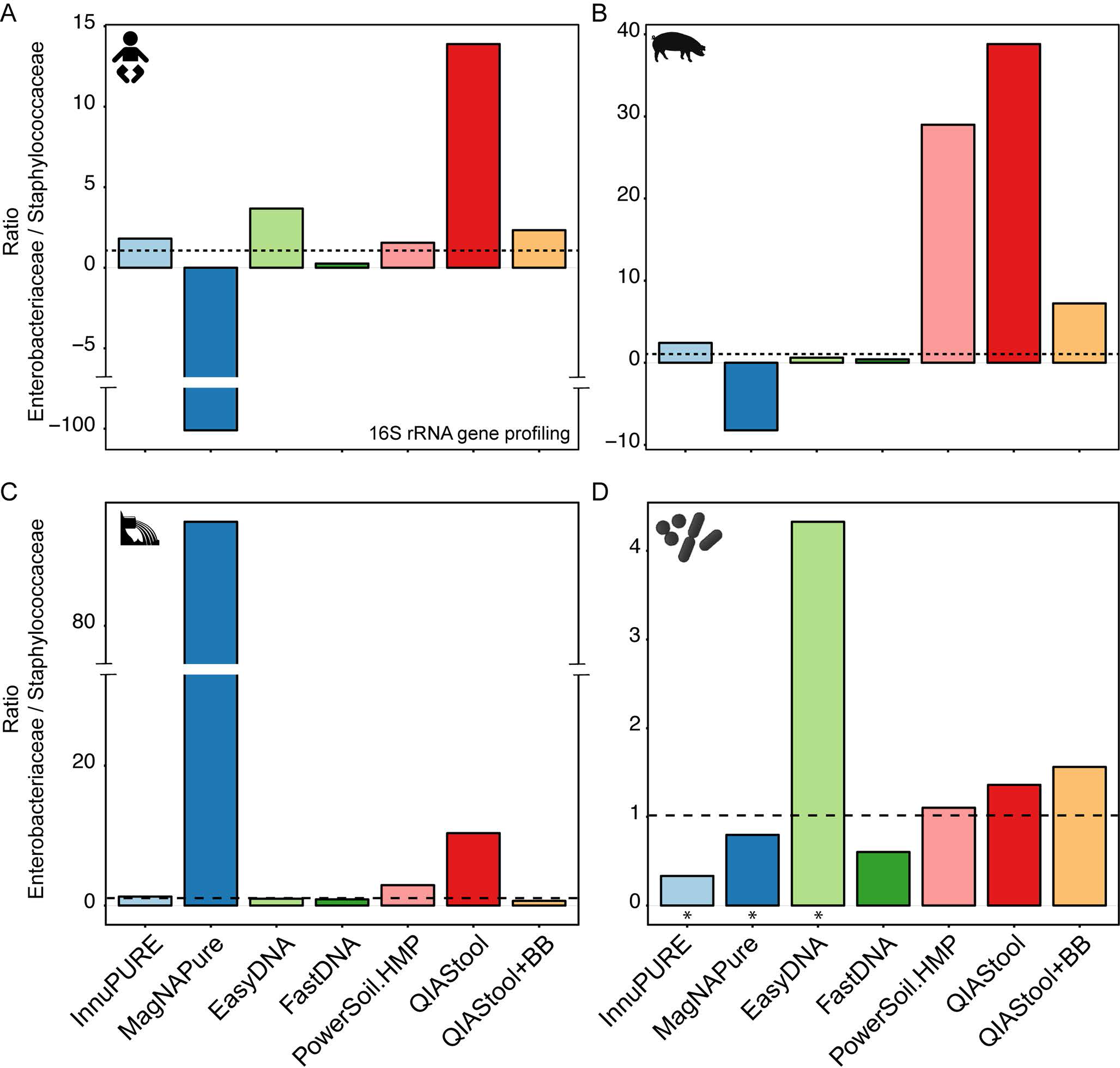
Detection of spiked bacteria. The human fecal (A), pig fecal (B), and hospital sewage (C) samples were spiked with a strain mix composed of *Salmonella enterica* serotype Typhimurium DT104 and *Staphylococcus aureus* ST398 in a CFU ratio of 1.02. The three sample matrices, as well as aliquots of the strain mix (D) were extracted using seven different DNA extraction methods. The two strains were detected by 16S rRNA gene profiling, and their ratios determined. For details, see Materials and Methods. An asterisk in (D) indicates that the values for the particular DNA extraction of the strain mix are based on single measurements. All other values are based on averages from duplicate or triplicate DNA extractions. The dashed line indicates the ratio of the strain mix based on CFU determinations. The x-axis scale is the same for all panels (A-D), and the y-axis scale specific for each sample type.

Overall, most DNA isolation methods exhibited a similar tendency across the three types of specimen. For example, for all three specimen types, the E/S ratio was higher using the QIAStool method, compared to the other methods (except MagNAPure for sewage). However, when the strain mix, composed of *S. enterica* and *S. aureus* only, was extracted using the seven DNA isolation procedures, their determined E/S ratio was in almost all cases similar to the expected ratio of 1.02, including the QIAStool method.

### Protocol modifications for increasing DNA concentration

One goal in genomics is to obtain a predicted pattern of microbial community composition that closely resembles the actual composition of microorganisms in a particular environment. Another challenge is to obtain sufficient DNA for metagenome sequencing. To address this aspect, we examined the effect of modifications to standard protocols on output DNA concentration (modifications are described in detail in the Supplemental Materials & Methods section). We chose the QIAStool method as a starting point, as we obtained DNA extracts using this method that were of high purity and stability (see Table S1A in the supplemental material). Another concern is processing time and costs for DNA isolation procedures, particularly for large-scale microbiome projects. The protocol of the QIAamp Fast DNA Stool Mini kit (QIAFast), a kit that became available at the time the present study was carried out, suggested reduced processing time compared to the QIAStool method. When we compared the QIAStool and QIAFast methods using metagenomic sequencing, we obtained a similar richness, diversity, and microbial community composition with these two methods (see Fig. S5 in the supplemental material). Furthermore, given that our previous results suggested that including a bead-beading step might result in a predicted community composition that was more similar to the community of known composition than without this step (Fig. 4), we included a bead-beating step and examined the effect of beads of differing types and cost (Table 1). We obtained a higher DNA concentration using pig feces and the QIAStool kit, when bead beating was applied and the double amount of volume after cell lysis was transferred (Fig. 5A). Similarly, for the QIAFast method, we obtained an on average 2.6-fold higher DNA concentration by including a bead beading step and transferring the double amount of volume after cell lysis, compared to DNA isolations without these modifications (Fig. 5A). Both, DNA purity and stability were in the expected range (see Table S3 in the supplemental material). Even though the DNA concentration was higher with these protocol modifications, the richness, diversity and community composition did not significantly differ when assessed by 16S rRNA gene profiling (Fig. 5A).

**FIG 5.**
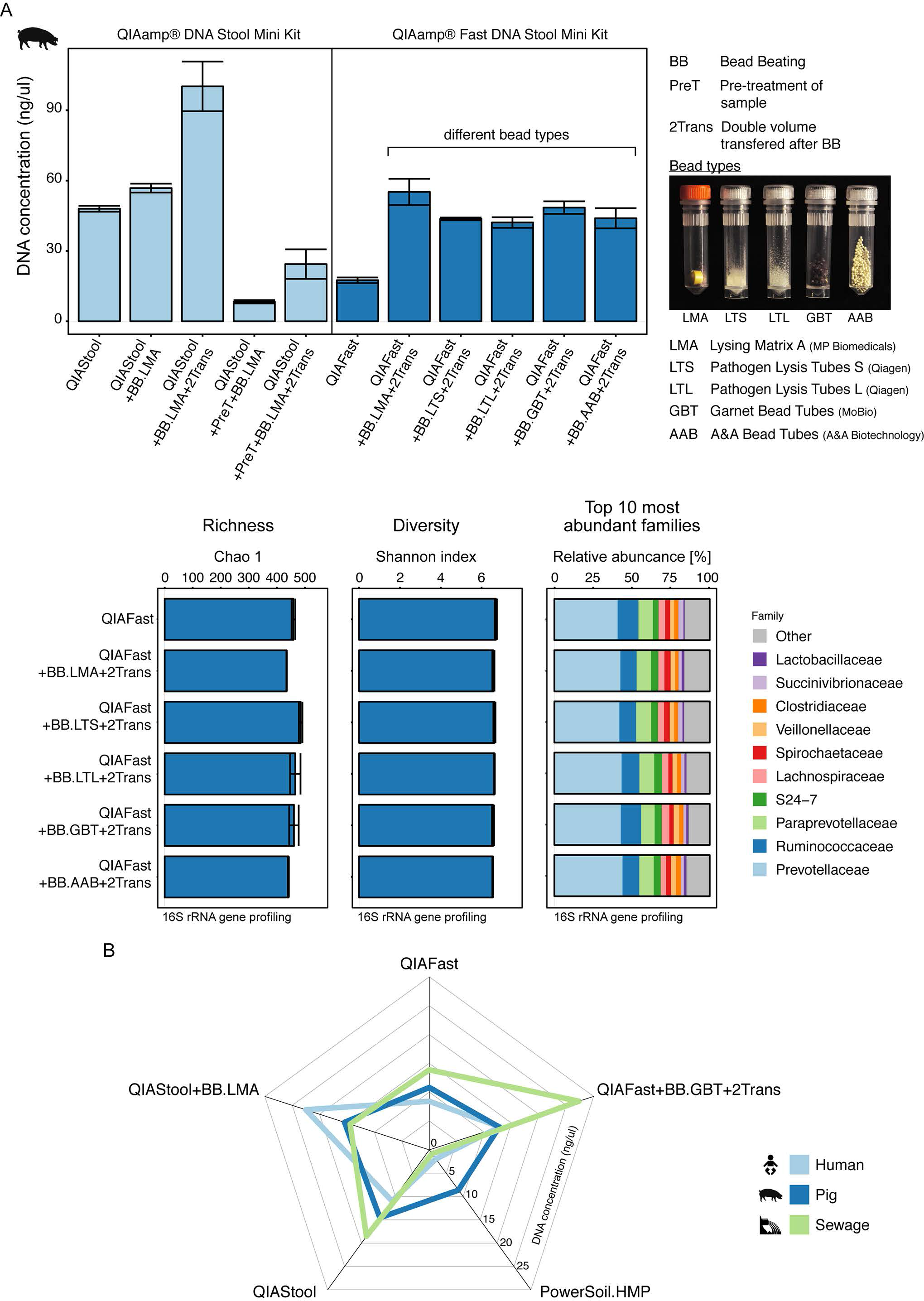
Effect of protocol modifications. A) Pig feces was extracted using standard as well as modified protocols based on the QIAamp^®^ DNA Stool Mini and QIAamp^®^ Fast DNA Stool Mini kits. The modifications included bead beading, pre-treatment of the sample, and transfer of the double amount of volume after cell lysis. In the bead-beating step, different bead types were examined (For details, see Materials and Methods, and Table 1). The alpha diversity (Chao 1 and Shannon index) was determined at OTU-level, and the microbial community composition examined at family-level based on 16S rRNA gene profiling. B) Selected standard and modified DNA extraction protocols were employed to extract DNA from human feces, pig feces, and sewage and their DNA concentration was displayed in a star plot. The values indicate the averages from duplicate extractions.

A particular DNA isolation method did not however lead to the highest DNA concentrations for each of the three types of specimen. Whereas the highest DNA concentration for sewage was achieved using the QIAFast+BB.GBT+2Trans method (27.30 ng/ul +/− 4.5 SEM), the highest DNA concentration for human feces was obtained using the QIAStool+BB.LMA method (22.50 ng/ul +/− 4.77 SEM) (Fig. 5B). For pig feces, the highest DNA concentrations were obtained using the QIAStool+BB.LMA (15.43 ng/ul +/−3.48 SEM) and QIAStool (14.57 ng/ul +/−3.62 SEM) methods. On average across the three types of specimen, the highest DNA concentrations were obtained using the QIAFast+BB.GBT+2Trans (17.66 ng/ul +/− 4.82 SEM) and QIAStool+BB.LMA (17.46 ng/ul +/− 2.54 SEM) methods.

## DISCUSSION

Genomics-based investigations of complex microbiomes greatly enhance our understanding about microbial community composition and function relevant to human, animal, and plant health, infectious diseases, environmental pollution, agriculture, and food safety. One current ambitious goal is to establish a global surveillance system for infectious agents and antimicrobial resistance based on next-generation DNA sequencing approaches (15). Given that infectious agents occupy various ecological habitats, DNA needs to be extracted from various types of specimen using standardized approaches in a time- and cost-efficient manner. It is advantageous, if a range of different specimens can be processed using the same standard operating procedure. In light of these considerations, we compared eight commercially available DNA isolation kits (a total of 16 protocols), and based on the findings developed an improved protocol using the QIAamp^®^ Fast DNA Stool Mini kit.

Overall, the amounts of DNA obtained from each DNA isolation method differed greatly, and there was no significant correlation between increasing DNA amount and increase in community diversity or richness. The taxonomic microbiome composition appeared to be dependent on both, the specimen and DNA isolation method. For example, the EasyDNA procedure preferentially extracted DNA from Gram-positive bacteria from the human feces and hospital sewage, while the FastDNA procedure preferentially extracted DNA from Gram-positive bacteria from pig feces. Methods that did not include a bead-beating or enzymatic treatment step generally extracted less DNA from Gram-positive bacteria. Furthermore, the results from our experiment that included the detection of spiked bacteria (Gram-negative and Gram-positive) suggests that quantification of distinct organisms from complex specimens is more challenging when the organisms are present at lower abundance levels. Inherent specimen properties may influence the DNA isolation efficiency leading to a biased pattern of microbial community composition. When using a particular procedure we found some similar abundance patterns of specific bacterial families between the three specimen types. However, we also observed several differences (e.g. Fig. 2 and Fig. 3). Hence, one cannot conclude that the DNA from a particular bacterial family will be extracted preferentially using one specific DNA isolation method across different types of specimens. This could be due to different inherent cellular properties of the taxa belonging to a specific family, affecting mechanical and enzymatic cell lysis. Moreover, the chemical and physical composition of the specimen could influence DNA isolation and downstream procedures. For example, it is well known that certain compounds, such as humic acid, polysaccharides, and bilirubin can affect PCR (16). Furthermore, fecal sample consistency, reflecting differences in water content and activity, can impact on microbial community composition (17).

Our observations from 16S rRNA gene profiling and metagenomics generally agreed, but the taxonomic patterns also exhibited some differences. One reason could be the known primer biases towards certain taxa in 16S rRNA gene based analysis (18). An additional reason could be differences in the composition of the reference databases used for the two sequence-based strategies. While 16S rRNA gene databases are composed of 16S rRNA gene sequences from a large diversity of taxa, the metagenomic sequence databases are based on whole and draft genome sequences from fewer and less diverse taxa. Both strategies complement each other, and efforts are ongoing in developing harmonized analytical workflows for sequence-based microbial community analysis.

Based on the insight gained in this study, we have developed an improved DNA isolation method based on the QIAamp^®^ Fast DNA Stool Mini kit. This procedure includes a bead beading step to obtain DNA from both, Gram-positive and Gram-negative taxa, and a step in which the double amount of cell lysate is transferred to the column to increase the DNA quantity. For aqueous sample types, like sewage, additional modifications are included, such as increasing the input amount and processing aliquots in parallel, as described in the SOP. While there was no single approach among the 16 procedures tested that appeared to completely resolve all challenges, we find the SOP based one the QIAamp^®^ Fast DNA Stool Mini kit useful for a number of reasons, including: 1) DNA extracts contained high amounts of DNA (sufficient to permitting PCR-free metagenomic sequencing) with high reproducibility 2) DNA extracts were of high quality in terms of DNA purity and stability, 3) DNA from both, Gram-positive and Gram-negative bacteria were reasonably well extracted (including Bifidobacteria), as determined by 16S rRNA amplicon profiling and metagenomic sequencing of spiked and un-spiked complex samples, 4) the method worked well for all examined sample types based on the DNA quality assessment and inferred microbiota composition, 5) the reagents and materials required were cheaper, and 6) the time needed for carrying out the DNA isolation was shorter, compared to several of the other procedures. A standard operating procedure for this DNA isolation method is available from <u>https://dx.doi.org/10.6084/m9.figshare.3475406</u>, and which can be used for different specimen types, and may be relevant to projects like EFFORT-against-AMR, COMPARE-Europe, the International Microbiome Initiative, and International Human Microbiome Standards.

In summary, our findings provide new insight into the effect of different specimen types and DNA isolation methods on DNA quantities and genomic-based inference of microbiome composition. We offer an optimized strategy for the DNA isolation for different sample types providing a representative insight into community composition, and which can be conducted in a time- and cost-efficient manner.

## MATERIAL AND METHODS

### Specimen Collection and Handling

Human fecal specimens were collected from a healthy individual. Pig fecal specimens were collected from animals at a conventional pig production farm in Denmark. Untreated sewage was collected from the sewage inlet of the Herlev hospital waste water treatment plant, Denmark. For details regarding sample handling and processing, see Supplemental Materials and Methods (Text S1).

### Spiking with strain mix

Subsequent to specimen collection, about half of the aliquots from the human, pig, and sewage were spiked with a representative of Gram-positive and Gram-negative bacteria, namely *Staphylococcus aureus* ST398 (strain S0385) and *Salmonella enterica* serotype Typhimurium DT104. For details regarding the preparation of the strain mix, see Supplemental Materials and Methods (Text S1).

### DNA isolation

In a first step, seven DNA isolation procedures were examined, namely: InnuPure^®^ C16, Analytic Jena AG (InnuPURE); MagNA Pure LC DNA isolation Kit III, Roche (MagNAPure); Easy-DNA™ gDNA Purification Kit, Invitrogen (EasyDNA); MP FastDNA™ Spin Kit, MP Biomedicals (FastDNA); PowerSoil^®^ DNA Isolation kit, MoBio (PowerSoil.HMP); QIAamp^®^ DNA Stool Mini Kit, Qiagen (QIAStool); QIAamp^®^ DNA Stool Mini Kit +Bead Beating, Qiagen (QIAStool+BB) (see Table 1, and details below). In a second step, a variety of modifications to two Qiagen kits were examined, namely the QIAamp^®^ DNA Stool Mini Kit (QIAStool), and QIAamp^®^ Fast DNA Stool Mini Kit (QIAFast). The standard operating procedure for an improved DNA isolation method (i.e. QIAamp Fast DNA Stool Modified, corresponding to QIAFast+BB.GBT+2Trans described here) can be found at https://dx.doi.org/10.6084/m9.figshare.3475406. For details regarding the individual DNA isolation procedures, see Supplemental Materials and Methods (Text S1).

### DNA quantitation and quality assessment

Subsequent to DNA isolation, the DNA was portioned into 10-μl aliquots to prevent repeated freeze-thawing cycles, and stored at −20°C. DNA concentrations were measured using Qubit^®^ dsDNA BR Assay Kit on a Qubit^®^ 2.0 Fluorometer (Invitrogen, Carlsbad, CA). As DNA extracts can contain contaminants, such as proteins and other organic molecules that can affect downstream procedures such as DNA amplifications in PCR, we determined the DNA purity by measuring the ratios of absorbance at 260/280 and 260/230, respectively, using a NanoDrop 1000 Spectrophotometer (Thermo Scientific, Pittsburgh, USA). DNA extracts with a 260/280 ratio between ~1.7 to ~ 2.0, and 260/230 ration between ~2.0 to ~2.2 are regarded as “pure”. The stability of the DNA in the extracts was determined by measuring the DNA concentration after 2 and 7 days incubation at 22°C. A decrease in DNA concentration over time can indicate the presence of DNases in the extract.

### 16S rRNA gene profiling

16S rRNA gene amplicon libraries were generated using a two-step protocol similar as described in Part # 15044223 Rev. B by Illumina. In a first PCR, the V4 region of the 16S rRNA genes were amplified using the universal primers (515f 5’- TGCCAGCAGCCGCGGTAATAC (19) and 806r 5’-GGACTACNNGGGTATCTAAT (20).

The samples were pooled in equal concentrations, and concentrated using ‘DNA clean and concentrator-5 kit’ (Zymo Research, Orange, CA). Paired-end 2 × 250 bp sequencing of barcoded amplicons was performed on a MiSeq machine running v2 chemistry (Illumina Inc., San Diego, CA, USA). The sequences were processed using the UPARSE pipeline (21) and a OTU x sample contingency table was created. Using QIIME1.8.0 (22), taxonomy was assigned with uclust using assign_taxonomy.py based on the Greengenes 13.8 reference database. Ecological diversity estimates and microbial community comparisons were performed using the relevant scripts provided by QIIME, phyloseq, and R (22–24). For details regarding the 16S rRNA gene-based microbial community analysis, see Supplemental Materials and Methods (Text S1), and the additional material provided through Figshare, https://figshare.com/projects/DNA_Isolation_Methodology_for_Microbiome_Genomics/14774.

### Metagenomics

A subset of the DNA extracts was subjected to metagenomic sequencing. The samples were prepared and sequenced following the Nextera XT DNA Library Preparation Guide for the MiSeq system Part # 15031942 Rev. D, using paired-end v2 2×250bp sequencing. The taxonomic microbiome compositions were determined through the use of the MGmapper pipeline (25). The MGmapper package is available for download at www.cbs.dtu.dk/public/MGmapper/. For details regarding the metagenomics-based microbial community analysis, see Supplemental Materials and Methods (Text S1).

### Differential abundance analysis

In order to test for the differential abundance of taxa that may drive the differences observed between the communities derived from the different DNA isolation procedures, we performed DESeq2 analyses. The read count tables from the 16S rRNA gene profiling and metagenomics sequence analysis, respectively, were aggregated to the family level in R (v. 3.2.3, 64bit) (24) We performed an analysis that allows for varied sequencing depth, similar as suggested previously (26), and carried out two-sided Wald tests as implemented in the DESeq2 (v. 1.10.1) package (27). The size factors were determined by DESeq2 from the read count tables. For details regarding the differential abundance analysis, see Supplemental Materials and Methods (Text S1).

### Quantification of strain mix

The samples that were spiked with the strain mix composed of *S. enterica* Typhimurium DT104 and *S. aureus* ST398 were extracted, sequenced, and analyzed together with the non-spiked samples. For each type of specimen and isolation method, the abundance of Enterobacteriaceae and Staphylococcaceae for 16S rRNA gene profiling and metagenomics, respectively, were determined. The ratios between Enterobacteriaceae and Staphylococcaceae was determined for each sample matrix and isolation method, and compared to the *S. enterica* Typhimurium DT104 / *S. aureus* ST398 ratio of CFU that were added to the original samples. For details regarding the quantification of the strain mix, see Supplemental Materials and Methods (Text S1).

### Ethics

The collection of human and pig fecal specimens as well as sewage was non-invasive, and were performed in accordance with the Declaration of Helsinki, and complied with Danish and European directives (86/609/EEC). The collection of specimens was conducted in accordance with the act on research ethics of heath research projects as administrated and confirmed by the National Committee on Health Research Ethics of Denmark (Region Hovedstaden), Journal nr. H-14013582.

### Accession numbers

The 16S rRNA gene sequences are available through the INSDC, such as from the European Nucleotide Archive (ENA) at the European Bioinformatics Institute (EBI) under accession number PRJEB12431, and the metagenomic sequences from ENA at EBI under accession number PRJEB14814.

## ACKNOWLEDGEMENTS

We thank Karin Vestberg (University of Copenhagen), Christina A. Svendsen (Technical University of Denmark) and Jacob D. Jensen (Technical University of Denmark) for technical assistance related to DNA sequencing. Marie S. Jensen (Technical University of Denmark) is acknowledged for the collection of pig fecal samples.

## AUTHOR CONTRIBUTIONS

B.E.K., L.B., F.M.A., and S.J.P. designed the research; B.E.K., L.B., O.L., and P.M. performed the research; B.E.K., L.B., O.L., P.M., A.P., F.M.A., and S.J.P. contributed analytic tools; B.E.K., L.B., O.L., P.M., and S.J.P. analyzed the data; B.E.K., L.B., and S.J.P. wrote the manuscript; and O.L., P.M., and A.P. edited the manuscript. All authors have read and approved the manuscript as submitted.

## SUPPLEMENTAL MATERIAL LEGENDS

**Text S1. Supplemental Materials and Methods.** Details regarding specimen collection and handling, spiking with the strain mix, DNA isolation, DNA quantitation and quality assessment, 16S rRNA gene profiling, metagenomics, differential abundance analysis, and quantification of the strain mix are described.

**FIG S1.**
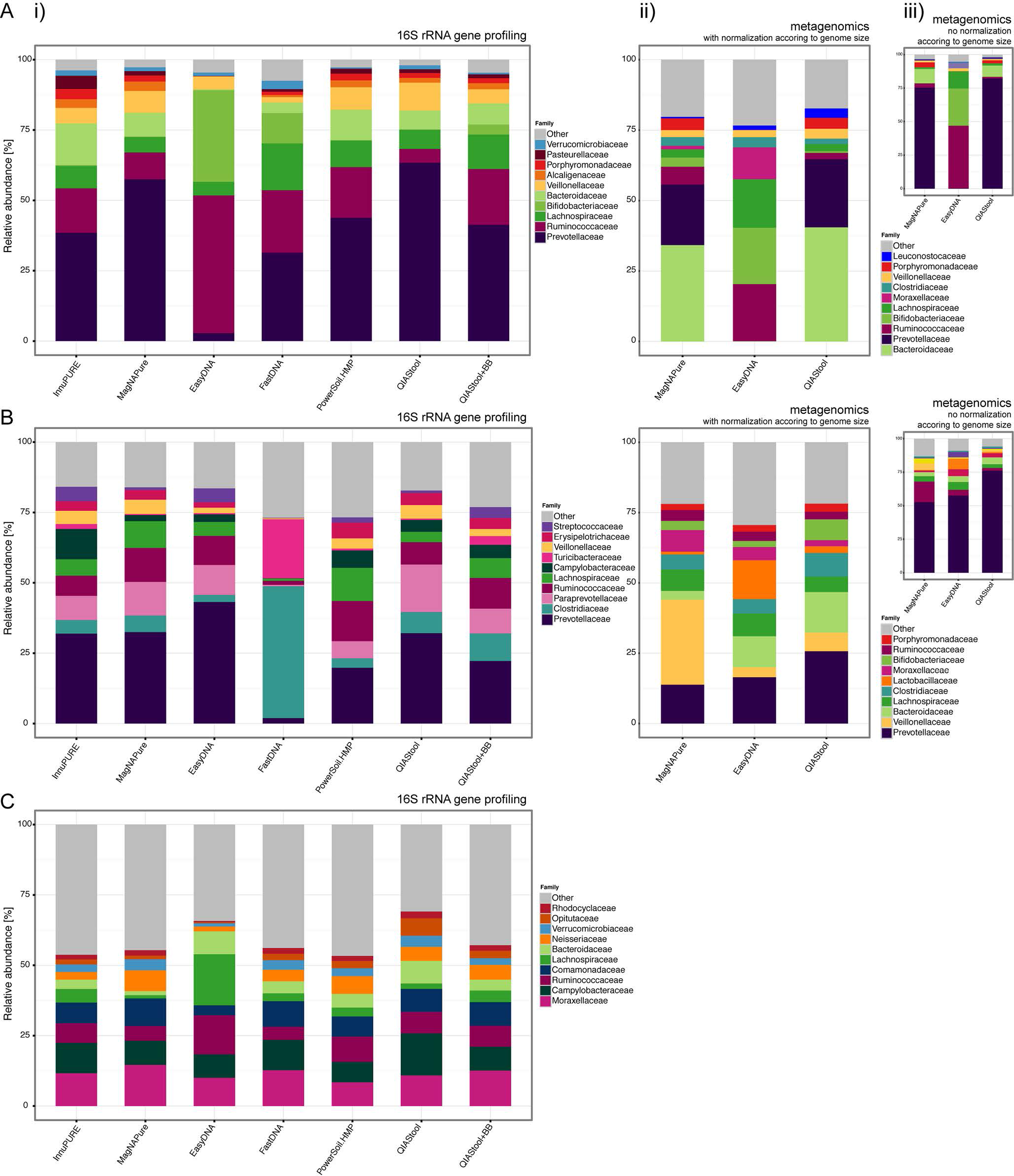
Microbial community composition. The Top 10 most abundant families for the human fecal (A), pig fecal (B), and hospital sewage (C) samples based on (i) 16S rRNA gene profiling, (ii) metagenomics analysis that include normalization based on reference genome size, and (iii) metagenomics analysis without normalization according to genome size. For details regarding sequence data analysis and normalization see Materials and Methods.

**FIG S2.**
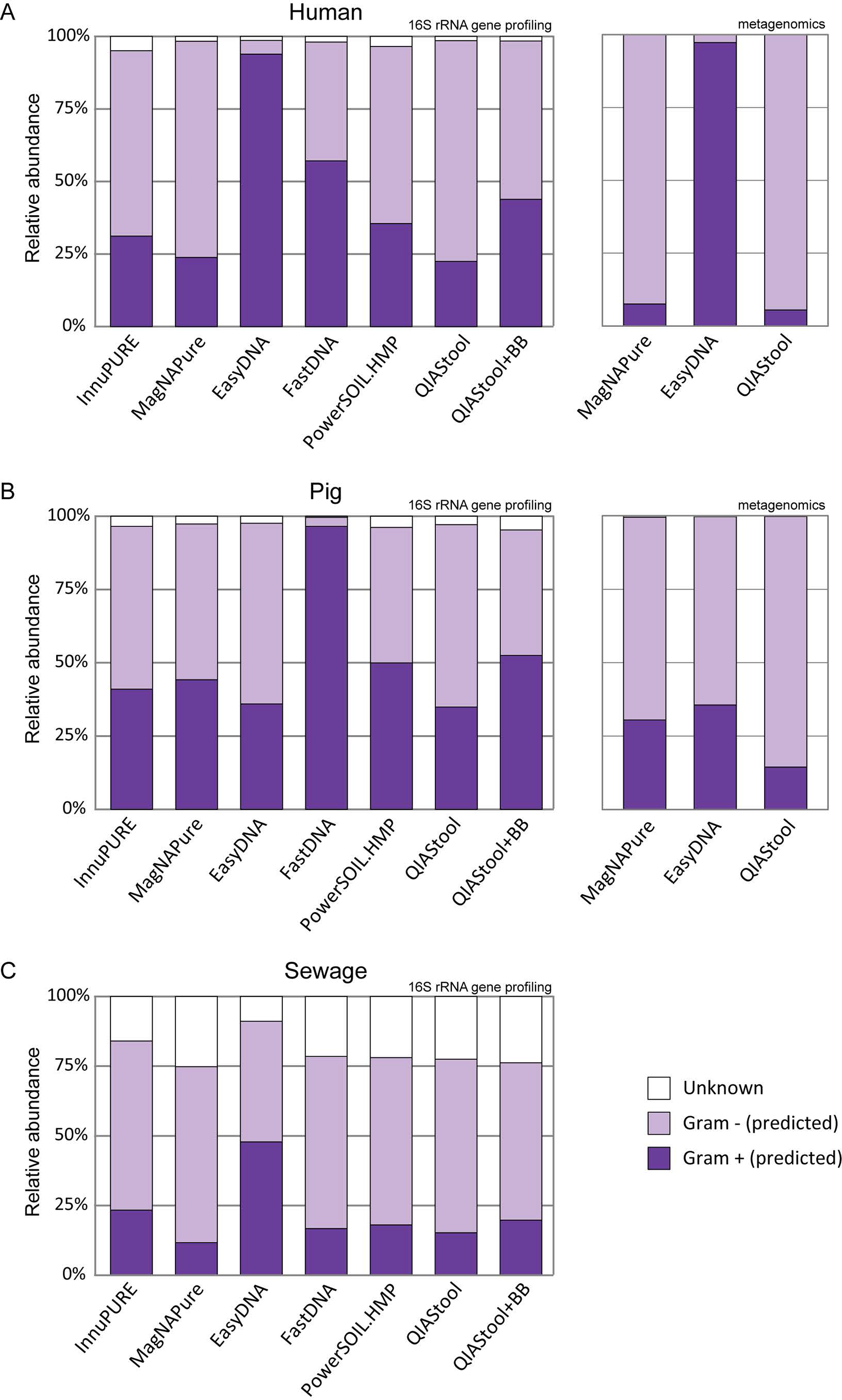
Microbial community composition based on predicted Gram-staining. Gram-positive and Gram-negative affiliations were assigned at the order-level based on information found in the literature. For some taxa the Gram-staining status was unknown.

**FIG S3.**
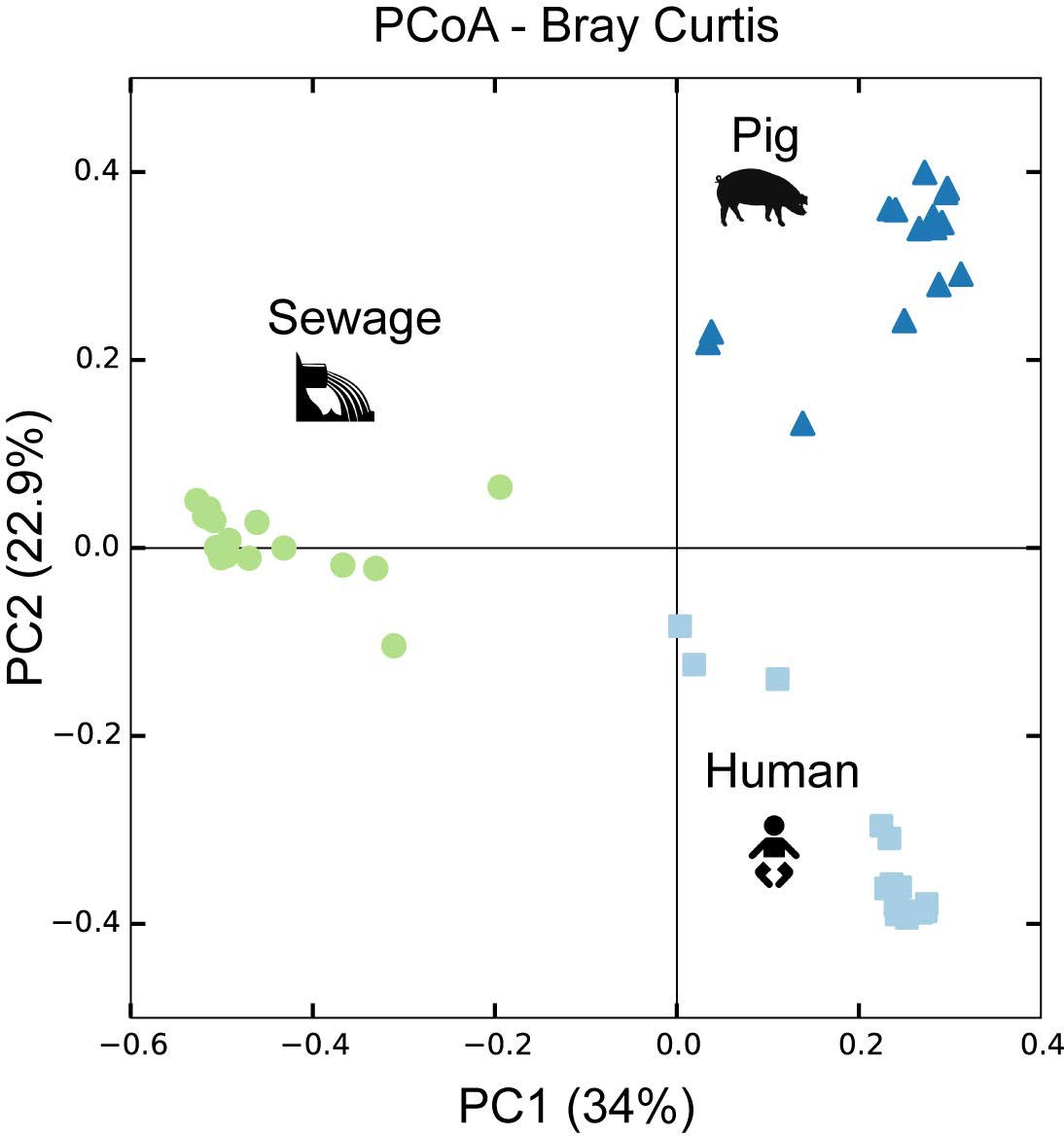
Microbial community dissimilarity. The dissimilarity between the microbiotas from the human, pig, and sewage samples was examined using Principal Coordinates Analysis of Bray-Curtis distances based on the 16S rRNA gene count data. For the PCoA Bray-Curtis ordination analysis only samples with a minimum of 800 reads were included. Additional results regarding community dissimilarity (based on Bray Curtis) and similarity (based on Jaccard similarity coefficient) within and between DNA extraction procedures across sample types as well as for a given sample type, are available through figshare: https://dx.doi.org/10.6084/m9.figshare.3814239.

**FIG S4.**
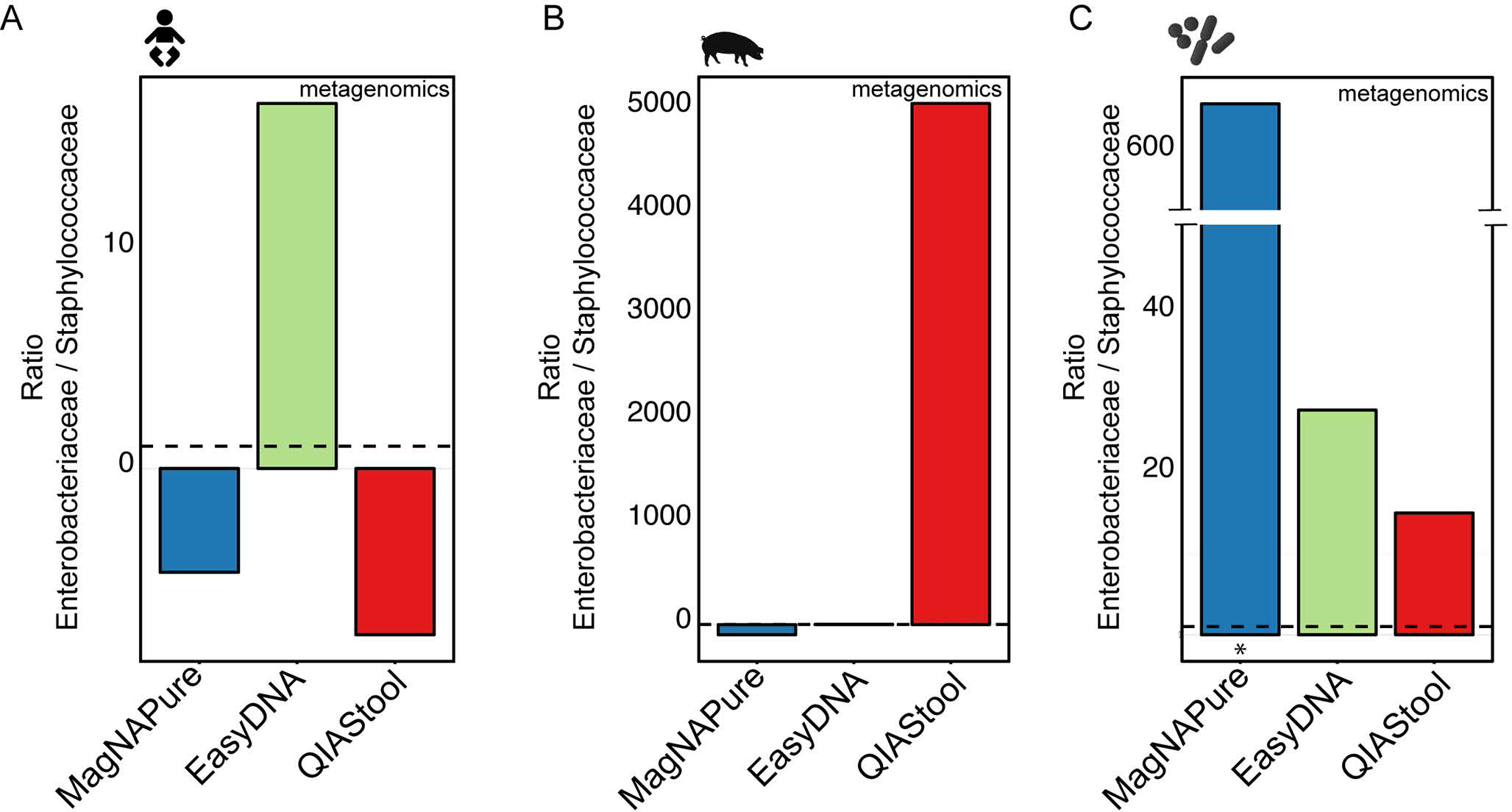
Detection of spiked bacteria using metagenomics. The human fecal (A), and pig fecal (B) samples were spiked with a strain mix composed of *Salmonella enterica* serotype Typhimurium DT104 and *Staphylococcus aureus* ST398 in a CFU ratio of 1.02. These two sample matrices, as well as aliquots of the strain mix (C) were extracted using three different DNA extraction methods. The two strains were detected by metagenomics analysis, and their ratios determined. For details, see Materials and Methods. An asterisk indicates that the values for the particular DNA extraction of the strain mix (D) are based on single measurements. All other values are based on averages from duplicate or triplicate measurements. The dashed line indicates the ratio of the strain mix based on CFU determinations.

**FIG S5.**
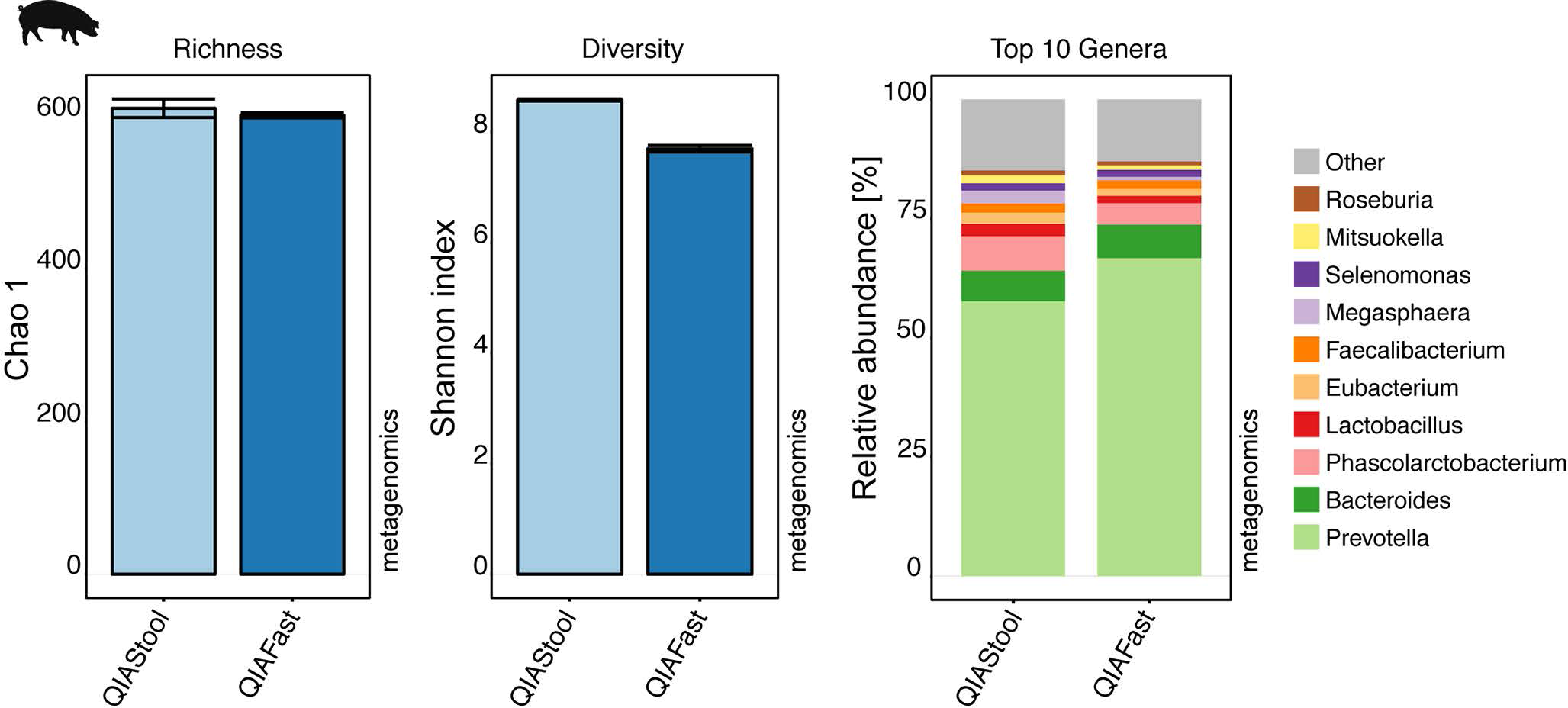
Comparison between QIAStool and QIAFast DNA extraction methods by metagenomics. Pig feces was extracted using the QIAamp^®^ DNA Stool Mini and QIAamp^®^ Fast DNA Stool Mini kits, and analyzed using metagenomics. The alpha diversity (Chao 1 and Shannon index) was determined at species-level. The microbial community composition was examined at genus-level and the relative abundance of the Top 10 most abundant taxa are shown here.

**Table S1. Comparison of DNA extraction methods.** (A) DNA concentration, purity, and stability, and (B) Microbiome richness and diversity.

**Table S2. Differential abundance of families.** (A) Human fecal microbial community, (B) Pig fecal microbial community, (C) Hospital sewage microbial community.

**Table S3. Comparison of DNA extraction methods.** DNA concentration, purity, and stability, for different DNA isolation procedures based on the QIAamp^®^ DNA Stool Mini and QIAamp^®^ Fast DNA Stool Mini Kits.

## SUPPLEMENTAL MATERIAL AND METHODS

### Specimen Collection and Handling

Human fecal specimens were collected from a healthy individual at three time points over a single day. The specimens were kept at 4°C, and transported to the laboratory within 24 hours. Upon arrival, the three samples were pooled and homogenized. For this study, fecal specimens from an infant were chosen, as infant fecal samples often contain a high proportion of Actinobacteria (e.g. Bifidobacteria), from which genomic DNA can be difficult to isolate. Pig fecal specimens were collected from animals at a conventional pig production farm in Denmark. Samples from individual animals were obtained directly after defecation, stored in a cooling box, and transported to the laboratory within four hours. Upon arrival, three random samples were pooled and homogenized. Untreated sewage was collected from the sewage inlet of the Herlev hospital waste water treatment plant, Denmark. Specimens were stored in a cooling box and transported to the laboratory within two hours. Upon arrival 24 × 40 ml sewage samples were sedimented for 10 minutes at 8000xg in an Eppendorf 5810R centrifuge. The sewage pellets were pooled and homogenized. For all three types of specimen (human feces, pig feces, sewage), the homogenized samples were separated into 0.5 g aliquots, respectively. A subset of aliquots for each specimen type was spiked with two bacterial strains (see details below). The individual sample aliquots with and without strain mix were stored at −80°C until further processing.

### Spiking with strain mix

Subsequent to specimen collection, about half of the aliquots from the human feces, pig feces, and sewage were spiked with a representative of Gram-positive and Gram-negative bacteria, namely *Staphylococcus aureus* ST398 (strain S0385) and *Salmonella enterica* serotype Typhimurium DT104. The strains were cultivated in Luria-Bertani (LB) broth at 37°C. Cells were harvested when the culture reached late exponential growth phase at OD _600_ ~0.9. The strain mix was prepared by mixing equal volumes of the bacterial cultures. To determine the number of cells of *S. aureus* ST398 and *Salmonella* Typhimurium DT104 in the two cultures, dilutions of these were plated on LB agar, the plates incubated overnight at 37°C, and colony forming units (CFU) determined the following day. The strain mix was added at about 5% of the volume of the aliquot, and the added cell numbers of *S. aureus* and *S. enterica* Thyphimurium were calculated based on the CFU determinations.

### DNA isolation

In a first step, seven DNA isolation procedures were examined, namely: InnuPure^®^ C16 (Analytic Jena AG), MagNA Pure LC DNA isolation Kit III (Roche), Easy-DNA™ gDNA Purification Kit (Invitrogen), MP FastDNA™ Spin Kit (MP Biomedicals), PowerSoil^®^ DNA Isolation kit (MoBio), QIAamp^®^ DNA Stool Mini Kit (Qiagen), QIAamp^®^ DNA Stool Mini Kit (Qiagen) +Bead Beating (see Table 1, and details below). These methods were selected because they are widely used and represent a variety of isolation procedures involving manual or automated DNA isolation, DNA separation using filter-columns or magnetic beads, chemical or mechanical lysis, and phenol/chloroform-based or non-chloroform based isolations. Bead-beating steps were performed in a Qiagen TissueLyser II if not stated otherwise, and centrifugation steps were carried out in an Ole Dich 157.MP Microcentrifuge (Denmark). DNA isolation was performed on duplicate or triplicate aliquots, dependent on specimen availability. One to two isolation controls were included at each round of isolation.

#### *InnuPure^®^ C16, Analytic Jena AG* (InnuPURE)

Automatic isolation with the InnuPURE-C16 robot using the InnuPURE Stool DNA Kit-IP-C16 according to the manufacture’s instructions. Prior to the automatic isolation, a lysis step was performed according to the protocol for lysis of bacterial DNA from stool samples using a SpeedMILL PLUS provided by the manufacturer. The cell disruption process was carried out two times for 30 sec at 50 Hz (50 s^−1^). The DNA was eluted in 100 ul of buffer supplied with the kit.

#### *MagNA Pure LC DNA isolation Kit III, Roche* (MagNAPure)

Automatic isolation with the MagNA Pure LC instrument using the DNA Isolation Kit III (Bacteria, Fungi) according to the manufacture’s instructions. The pre-isolation step for stool samples described in the protocol was performed before transferring the samples to the MagNA Pure LC. The protocol states a starting amount of a peanut-size sample, and in order to ensure consistency across isolations a starting amount of 0.25 g was chosen. The DNA was eluted in 100 ul of buffer supplied with the kit.

#### *Easy-DNA™ gDNA Purification Kit, Invitrogen* (EasyDNA)

The DNA isolation was performed according to the manufacturer’s instructions with minor modifications. Pretreatment of the samples were performed following the protocol for small amounts of cells, tissues, or plant leaves. Initially, 0.25 g sample aliquots were resuspended in 1.5 ml 0.9% NaCl, respectively. The samples were centrifuged at 600xg for 3 minutes. The supernatant was transferred to new tubes and centrifuged at 8000xg for 10 minutes. After centrifugation, the supernatant was discarded and the pellet was resuspended in 200 μl PBS. 30 μl lysozyme (10 mg/ml) and 15 μl lysostaphin (10 mg/ml) were added and the samples were incubated at 37°C for 20 minutes shaking at 550 rpm, before adding 30 μl 10% SDS. The final pretreatment step included the addition of 15 μl proteinase K (20 mg/ml) and incubation at 37°C for 20 minutes. The final step in the isolation protocol was prolonged to an incubation for 1.5 (instead of 0.5) hours at 37°C. The DNA was eluted in 100 ul of buffer supplied with the kit.

#### *MP FastDNA™ Spin Kit, MP Biomedicals* (FastDNA)

The DNA isolation was performed according to the manufacturer’s instructions with minor modifications. A centrifugation step at 3000xg for 2 minutes was included to ensure proper settling of the silica matrix. The protocol suggested eluting the DNA in 50-100 μl DNase/Pyrogen-Free water, and here the DNA was eluted in 100 μl.

#### *PowerSoil^®^ DNA Isolation kit, MoBio Laboratories Inc.* (PowerSoil.HMP)

The DNA isolation was performed according to the protocol employed in the Human Microbiome Project (HMP Protocol # 07-001 version 12), with a minor modification to the initial protocol step. The HMP protocol states to resuspend 2 ml fecal sample in 5 ml MoBio lysis buffer. Here, we resuspended 0.5 g sample in 1.25 ml MoBio lysis buffer (i.e. same ratio). Subsequently, the samples were centrifuged according to the HMP protocol and 1 ml of supernatant transferred to a garnet bead tube containing 0.75 ml MoBio buffer. The samples were heated at 65°C for 10 minutes followed by an additional heating step at 95°C for 10 minutes. The samples were processed further according to the HMP protocol including the modification at step 12, where the centrifugation step was prolonged to 2 minutes. The DNA was eluted in 100 ul of buffer supplied with the kit.

#### *QIAamp^®^ DNA Stool Mini Kit, Qiagen* (QIAStool)

The DNA isolation was performed according to the manufacturer’s protocol for isolation of DNA from stool for pathogen detection with minor modifications. In the lysis step, the samples were first heated at 70°C for 5 minutes and subsequently at 95°C for 5 minutes. The DNA was eluted in 100 μl elution buffer.

#### *QIAamp^®^ DNA Stool Mini Kit, Qiagen* +Bead Beating (QIAStool+BB)

The DNA isolation was performed according to the manufacturer’s protocol for isolation of DNA from stool for pathogen detection with minor modifications that included a bead-beating step. Sample aliquots of 0.2 g were mixed with 1.4 ml ASL buffer, respectively, and added to Lysing matrix A bead beating tubes (MP Biomedicals) and were briefly homogenized. The samples were treated in a Qiagen TissueLyser II at 30 f/s (Hz) three times for 30 seconds, with placement of the samples on ice in between bead beating steps. Subsequently, the samples were heated at 95°C for 15 minutes. The remaining steps were carried out according to the manufacturer’s recommendations, and the DNA eluted in 100 ul elution buffer.

In a second step, a variety of modifications to two Qiagen kits were examined, namely the QIAamp^®^ DNA Stool Mini Kit (QIAStool), and QIAamp^®^ Fast DNA Stool Mini Kit (QIAFast). The latter was released to the market during the course of this study. The main difference between the QIAStool and the QIAFast kits relies in the way inhibitor compounds are being removed. In the QIAStool kit, InhibitEX tablets are being dissolved in the samples that are adsorbing the inhibitors and together are removed via centrifugation. The QIAFast kit contains an InhibitEX buffer to remove inhibitor compounds, and no tablets are required.

#### *QIAamp^®^ DNA Stool Mini Kit, Qiagen* and Modifications

Five different protocols based on the QIAamp^®^ DNA Stool Mini Kit were examined. i) QIAStool: see above, ii) QIAStool+BB.LMA: QIAStool+BB procedure using Lysing matrix A tubes (see above), iii) QIAStool+BB.LMA+2Trans: QIAStool+BB procedure using Lysing matrix A tubes (see above) with modifications. To reduce the loss of sample, the double amount of supernatant was transferred to proteinase K (i.e. 400 μl instead of 200 μl). The volumes of Proteinase K, buffer AL and ethanol were doubled, respectively. Due to the increased volume, the passing of the sample through the spin columns is performed in two centrifugation steps. The DNA was washed twice before elution in 100 μl elution buffer. iv) QIAStool+PreT+BB.LMA: QIAStool+BB procedure using Lysing matrix A tubes (see above) with modifications. An increased starting sample amount was used and pre-treated. 0.5g of sample was mixed with 1.5 ml 0.9% NaCl solution. After homogenization by vortexing, the samples were centrifuged at 600 x g for 3 minutes to settle large particles. The supernatant was centrifuged at 8000 x g for 10 minutes to pellet microbial cells. The pellet was resuspended in 200 μl PBS and transferred to Lysing Matrix A bead beating tubes. v) QIAStool+PreT+BB.LMA+2Trans: QIAStool+BB procedure using Lysing matrix A tubes (see above) with modifications described in iii) and iv).

#### *QIAamp^®^ Fast DNA Stool Mini Kit, Qiagen* and Modifications

Six different protocols (i-vi) based on the QIAamp^®^ Fast DNA Stool Mini Kit were examined using five different bead types (ii-vi). i) QIAFast: The DNA isolation was performed according to the manufacturer’s protocol for isolation of DNA from stool for pathogen detection with minor modifications. In the lysis-step, the samples were first heated at 70°C for 5 minutes and subsequently at 95°C for 5 minutes. The DNA was eluted in 100 μl elution buffer for 2 minutes. ii) QIAFast+BB.LMA+2Trans: The DNA isolation was performed according to the manufacturer’s protocol for isolation of DNA from stool for pathogen detection with minor modifications that included a bead-beating step. Sample aliquots of 0.2 g were mixed with 1 ml InhibitEX buffer, respectively, and added to Lysing matrix A bead beating tubes (MP Biomedicals) and are briefly homogenized. The samples are treated in a Qiagen TissueLyser II at 30 f/s (Hz) three times for 30 seconds, with placement of the samples on ice in between bead beating steps. Subsequently, the samples are heated at 95°C for 7 minutes. Similar to the modifications described above, following the bead-beating and heating steps, the double amount of supernatant was transferred to proteinase K (i.e. 400 μl instead of 200 μl). The volumes of proteinase K, Buffer AL and ethanol were also doubled. The passing of the sample through the filter columns were subsequently carried out in two centrifugation steps rather than one, to accommodate the increased sample volume. The remaining steps were carried out according to the manufacturer’s recommendations, and the DNA eluted in 100 ul elution buffer. A laboratory protocol for this procedure can be found at <u>https://dx.doi.org/10.6084/m9.figshare.3475406</u>. iii) QIAFast+BB.LTS+2Trans: Same procedure as described in ii) with Pathogen Lysis Tubes S (Qiagen). iv) QIAFast+BB.LTL+2Trans: Same procedure as described in ii) with Pathogen Lysis Tubes L (Qiagen). v) QIAFast+BB.GBT+2Trans: Same procedure as described in ii) with Garnet Bead Tubes (MoBio). vi) QIAFast+BB.AAB+2Trans: Same procedure as described in ii) with A&A Bead Tubes (A&A Biotechnology, Gdynia, Poland).

Together, the evaluation and improvements of DNA isolation methods were carried out in a step-wise approach. In the first step, seven DNA extraction kits were evaluated using human feces, pig feces, and hospital sewage (Figures 1–4, Supplemental Figures S1–S4, and Supplemental Tables S1+S2). The standard and modified procedures based on the QIAStool and QIAFast methods were tested using a second set of pig fecal samples (Figure 5A, and Supplemental Figure S5, and Supplemental Table S3). Upon evaluation of the different DNA isolation methods, promising procedures were selected and examined using a new set of human feces, pig feces, and hospital sewage (Figure 5B).

### DNA quantitation and quality assessment

Subsequent to DNA isolation, the DNA was portioned into 10 μl aliquots to prevent repeated freeze-thawing cycles, and stored at −20°C. DNA concentrations were measured using Qubit^®^ dsDNA BR Assay Kit on a Qubit^®^ 2.0 Fluorometer (Invitrogen, Carlsbad, CA). As DNA extracts can contain contaminants, such as proteins or other organic molecules that can affect downstream procedures such as DNA amplifications in PCR, we determined the DNA purity by measuring the ratios of absorbance at 260/280 and 260/230, respectively, using a NanoDrop 1000 Spectrophotometer (Thermo Scientific, Pittsburgh, USA). DNA extracts with a 260/280 ratio between ~1.7 to ~ 2.0, and 260/230 ration between ~2.0 to ~2.2 are regarded as “pure”. The stability of the DNA in the extracts was determined by measuring the DNA concentration after 2 and 7 days incubation at 22°C. A decrease in DNA concentration over time can indicate the presence of DNases in the extract.

### 16S rRNA gene profiling

16S rRNA amplicon libraries were generated using a two-step protocol similar as described in Part # 15044223 Rev. B by Illumina (<u>http://www.illumina.com/con</u> <u>tent/dam/illumina-support/documents/documentation/chemistry_documenta tion/16s/16s-metagenomic-library-prep-guide-15044223-b.pdf</u>). In a first PCR, the V4 region of the 16S rRNA genes were amplified using the universal primers (515f 5’-TGCCAGCAGCCGCGGTAATAC (1) and 806r 5’-GGACTACNNGGGTATCTAAT (2). Each 20-μl PCR reaction contained 2 μl of 10 × AccuPrime PCR Buffer II (15mM MgCl2, Invitrogen), 1 μl (10 μM) of the primers, 0.12 μl AccuPrime Taq DNA polymerase (2 units/μl, Invitrogen), 1 μl template DNA and 14.88 μl ddH2O. PCR conditions: denaturation at 94°C for 2 min; 30 cycles at 94°C for 20 s, 56°C for 20 s, 68°C for 30 s; followed by 68°C for 5 min, and 3 min at 70 °C. Subsequently, the PCR products were placed on ice to prevent hybridization between PCR products and nonspecific amplicons. Samples were quantified using Quant-iT™ PicoGreen^®^ dsDNA Assay Kit (Invitrogen, Carlsbad, CA) on a Lightcycler 96 (Roche, Mannheim, Germany) and adjusted to equal concentrations. In the second PCR the same conditions were used as in the first round, except the PCR was reduced to 15 cycles and the primers had a unique adaptor/linker/index sequence per sample (3). The PCR products were purified using Agencourt AmPure XP beads (Beckman Coulter Inc, A63881), and concentrations were measured using Quant-iT™ PicoGreen^®^ dsDNA Assay Kit on a Lightcycler 96. The samples were pooled in equal concentrations, and concentrated using ‘DNA clean and concentrator-5 kit’ (Zymo Research, Orange, CA). Paired-end 2 × 250 bp sequencing of barcoded amplicons was performed on a MiSeq machine running v2 chemistry (Illumina Inc., San Diego, CA, USA) at University of Copenhagen, Section of Microbiology.

The primer sequences were trimmed, quality filtering performed, and paired sequences assembled using the UPARSE pipeline <u>http://drive5.com/usearch/manual/uparse_pipeline.html</u> (4). Low quality reads were removed with a maximum expected error threshold of 0.5 (maxee). Sequences were barcoded and pooled before dereplication and removal of duplicates (-minseqlength 64). Prior to clustering of the OTUs the dereplicated reads were sorted according to abundance, and singeltons where removed (<u>http://drive5.com/usearch/manual/upp_readprep.html</u>). Chimera filtering was performed using UCHIME (5) with rdp_gold.fa as reference database. The reads were mapped back to OTUs, including singletons, at a 97% identity level and an OTU-table was generated using uc2otutab.py. Using QIIME1.8.0 (6), taxonomy was assigned with uclust using assign_taxonomy.py based on the Greengenes 13.8 reference database. The average number of reads per sample was 192965, and the read length was between 186-251 bp. The average number of reads in the isolation controls was 34063 and the majority of these reads were affiliated with the two strains used for spiking (Enterobacteriaceae and Staphylococcaceae), as well as dominant taxa that were present in the complex samples, such as Ruminococcaceae, Prevotellaceae, and Bacteroidales. Ecological diversity estimates and microbial community comparisons were performed using the relevant scripts provided by QIIME, phyloseq, and R (6–8). For the estimation of bacterial diversity and richness (Fig. 1C, and Fig. 5A), and principal coordinate analysis (Fig. 2A-C, and Fig. S3) the samples were rarefied to 800 reads per sample. The abundance of Gram-positive and Gram-negative bacteria was predicted at order levels based on information from the literature. For some bacteria (mainly Firmicutes), the Gram status could not be assigned at this level, and for those the family level was used instead.

### Metagenomics

A subset of thirty-nine DNA extracts was subjected to metagenomic sequencing. The samples were prepared and sequenced following the Nextera XT DNA Library Preparation Guide for the MiSeq system, Part # 15031942 Rev. D (<u>http://support.illumina.com/content/dam/illumina-support/documents/documentation/chemistry_documentation/samplepreps_nextera/nextera-xt/nextera-xt-library-prep-guide-15031942-01.pdf</u>), using paired-end v2 2×250bp sequencing. The taxonomic microbiome compositions were determined through the use of the MGmapper pipeline (9). The MGmapper package is available for download at <u>www.cbs.dtu.dk/public/MGmapper/</u>. The analysis consisted of three main steps: i) Pre-processing and quality trimming of raw reads, ii) Mapping of reads to reference sequence databases, and iii) Analysis of read count data. In the first step, cutadapt (10) was employed for adapter sequence removal, trimming of low-quality bases from the ends of the reads (-q 30), and removal of reads that were shorter than 30 bp. In a second step, the remaining paired-end reads were mapped in chain-mode to four databases: 1. complete bacterial genomes, 2. draft bacterial genomes, 3. MetaHit Assembly (<u>http://www.sanger.ac.uk/resources/downloads/bacteria/metahit/</u>, July 2014), and 4. Human Microbiome assemby (<u>http://www.hmpdacc.org/resources/data_browser.php</u>, July 2014) using the BWA-MEM algorithm (<u>http://bio-bwa.sourceforge.net</u>). For the analysis in the present study only the reads mapping to the two primary bacterial databases (complete and draft bacterial genomes) were considered. These two databases were composed of 2685 complete and 22224 draft bacterial and archaeal genomes obtained from Genbank on July 2014 and December 2014, respectively. The order by which the databases are specified in chain-mode is important, as reads that exhibit a significant hit to the previous reference database are removed before mapping to the next database. Samtools (10) was used to remove singletons and all reads that did not map as pairs. An alignment of a read pair with a region in a genome was considered a hit only if the sum of the alignment scores (SAS) was higher than any SAS values from other hits in the database. In the third step, the alignments were filtered based on the Fraction of Matches+Mismatches (FMM) threshold, i.e. the fraction of a read that should align. Here, the default FMM threshold of 80% was used. From 96 155 142 raw read pairs, 7 567 574 read pairs mapped genomes in the two reference databases. The final read count table was composed of 9436 bacterial and archaeal reference strains with an average of 69952 mapped read pairs per sample. For each sample, the read counts were normalized according to the genome length of the respective genomes in the database, and for sequencing depth using total sum scaling.

### Differential abundance analysis

In order to test for the differential abundance of taxa that may drive the differences observed between the communities derived from the different DNA isolation procedures, we performed DESeq2 analyses. The (unnormalized) read count tables from the 16S rRNA gene profiling and metagenomics sequence analysis, respectively, were aggregated to the family level in R (v. 3.2.3, 64bit) (8). We performed an analysis that allows for varied sequencing depth, similar as suggested previously (12), and carried out two-sided Wald tests as implemented in the DESeq2 package (v. 1.10.1) (13). The size factors were determined by DESeq2 from the read count tables. An example for such an analysis is available from <u>https://dx.doi.org/10.6084/m9.figshare.3811251</u> (available as .Rmd, .html, and .pdf).

When testing the effect of added strain mix, we included the samples to which the strain mix was added as well as the corresponding samples to which no strain mix was added and accounted for DNA isolation method and sample matrix type. When testing the effects of the DNA isolation method, we analyzed the data from the three types of fecal specimen separately and extracted results from all two-wise comparisons of DNA isolation methods. For each DESeq2 test, p-values were adjusted for the false-discovery rate (FDR) using the Benjamini-Hochberg procedure (14). As recommended by DESeq2, comparisons with an FDR below 0.1 were considered significant. For the visualization of the data, the read count data were variance-stabilized using the DESeq2 regularized log (rlog) transformation. This transformation also accounts for sequencing depth differences, allowing inter-sample comparisons of taxa.

### Quantification of strain mix

The samples that were spiked with the strain mix composed of *S. enterica* Typhimurium DT104 and *S. aureus* ST398 were extracted, sequenced, and analyzed together with the non-spiked samples. For each type of specimen and isolation method, the relative abundance of Enterobacteriaceae and Staphylococcaceae for 16S rRNA gene profiling and metagenomics, respectively, were determined. Our differential abundance analysis using DESeq2 confirmed, that these two strains were present in significantly higher abundance in the spiked samples than in the not spiked samples for 16S rRNA gene profiling: Enterobacteriaceae adjusted P-value 3.08^−30^ and Staphylococcaceae adjusted P-value 2.13^− 10^; and for metagenomics: Enterobacteriaceae adjusted P-value 1.74^−77^ and Staphylococcaceae adjusted P-value 1.07^−4^. The average relative abundance values from the samples without added strain mix were subtracted from the corresponding samples to which the strain mix was added. Subsequently, the 16S rRNA gene copy numbers of the two added strains were taken into account with 5 for *S. aureus* and 7 for *S. enterica* (for 16S rRNA gene profiling). The ratios between Enterobacteriaceae and Staphylococcaceae were determined for each sample matrix and isolation method, and compared to the *S. enterica* Typhimurium DT104 / *S. aureus* ST398 ratio of CFU that were added to the original samples.

